# SPDI: Data Model for Variants and Applications at NCBI

**DOI:** 10.1101/537449

**Authors:** J. Bradley Holmes, Eric Moyer, Lon Phan, Donna Maglott, Brandi L. Kattman

**Affiliations:** National Center for Biotechnology Information (NCBI), National Library of Medicine (NLM), National Institutes of Health (NIH), 8600 Rockville Pike, Bethesda, MD 20894, USA

## Abstract

**Motivation:** Normalizing diverse representations of sequence variants is critical to the elucidation of the genetic basis of disease and biological function. NCBI has long wrestled with integrating data from multiple submitters to build databases such as dbSNP and ClinVar. Inconsistent representation of variants among variant callers, local databases, and tools results in discrepancies and duplications that complicate analysis. Current tools are not robust enough to manage variants in different formats and different reference sequence coordinates.

**Results:** The SPDI (pronounced “speedy”) data model defines variants as a sequence of 4 operations: start at the boundary before the first position in the sequence **S**, advance **P** positions, delete **D** positions, then insert the sequence in the string **I**, giving the data model its name, SPDI. The SPDI model can thus be applied to both nucleotide and protein variants, but the services discussed here are limited to the nucleotide. Current services convert representations between HGVS, VCF, and SPDI and provide two forms of normalization. The first, based on the NCBI Variant Overprecision Correction Algorithm, returns a unique, normalized representation termed the “Contextual Allele” for any input. The SPDI name, with its four operations, defines exactly the reference subsequence potentially affected by the variant, even in low complexity regions such as homopolymer and dinucleotide sequence repeats. The second level of normalization depends on alignment dataset (ADS). SPDI services perform remapping (AKA lift-over) of variants from the input reference sequence to return a list of all equivalent Contextual Alleles based on the transcript or genomic sequences that were aligned. One of these contextual alleles is selected to represent all, usually, that based on the latest genomic assembly such as GRCh38 and is designated as the unique “Canonical Allele”. ADS includes alignments between non-assembly RefSeq sequences (prefixed NM, NR, NG), as well inter- and intra-assembly-associated genomic sequences (NCs, NTs, and NWs) and this allows for robust remapping and normalization of variants across sequences and assembly versions.

**Availability and implementation:** The SPDI services are available for open access at: https://api.ncbi.nlm.nih.gov/variation/v0/

## Introduction

Understanding genetic variants and the relationships between phenotype and genotype is critical for genetic research and for the application of precision medicine [1]. Within the next decade, millions of genomes will be sequenced and billions of variants will be called from them to provide the data to determine the genetic contribution to diseases, cancers, and responses to drugs and treatments^1^. Crucial to variant analysis is the normalizing of representations from disparate callers and formats to compare both to user data sets as well as annotations cataloged in public databases such as dbSNP and ClinVar [3]. The normalization step can be challenging because any variant can be described using different formats (VCF, HGVS, and other identifiers) and different reference sequence types (mRNA, protein, and genomic) and assembly versions (hg18, hg19, NCBI36, GRCh37, or GRCh38). Previously, NCBI’s dbSNP, a database of small genetic variations, addressed this problem by aggregating variant data from disparate submitters by mapping the variant to the genome using defined sequences flanking the variant site. All novel and existing variant flanking sequences were mapped to the latest genomic assembly by BLAST [4]. Variants of the same type (SNV, deletion, *etc*.) and that mapped to the same genomic locations were assigned a unique dbSNP Reference SNP cluster ID (RSID) to create a non-redundant set of reference variants (RefSNP). The latest dbSNP build, 151 (April 2018 release), has over 650 million human RefSNPs that were created from over 2 billion submitted SNP (SubSNP). However, the algorithm employed by dbSNP was overly precise for variants in low complexity regions where reported variant position can be ambiguous (Figure 3). RefSNP clustering artifacts also resulted from imperfect alignments, errors in submitted flanking sequence, and ambiguous mapping in duplicated or repetitive genomic regions. Another roadblock to variant normalization has been different reporting formats commonly used by the community. The Human Genome Variation Society (HGVS) format is a recommended standard for describing variation in the literature [5]. The genomic community often reports variants in VCF format [6] with asserted locations anchored on a reference sequence. Tan et al. [7] have developed a variant normalization method that is left aligned and parsimonious for VCF format and a variant tool that combine and compare variants from two VCF files is available [8]. However, these tools are specific to VCF and do not work across other variant nomenclature formats such as HGVS. Furthermore, they lack an integrated remapping (lift-over) function to normalize variants reported on different reference sequences (mRNA, genomic loci, and contigs) and assembly versions. To address these issues, we developed the SPDI data model to improve variant representation along with algorithms and services to improve variant normalization. These services are available via a web API (https://api.ncbi.nlm.nih.gov/variation/v0) to aid users in variant analysis and interpretation, and now underpin the dbSNP and ClinVar dataflows to aggregate submissions and reannotate variants for regular releases.

## Methods

### Variant Model

For nucleotides, SPDI represents all variants as a sequence of 4 operations (Table 1): start at the boundary before the first nucleotide in the sequence S, advance P nucleotides, delete D nucleotides, then insert the nucleotides in the string I. This model has four parameters:

**Table 1:**
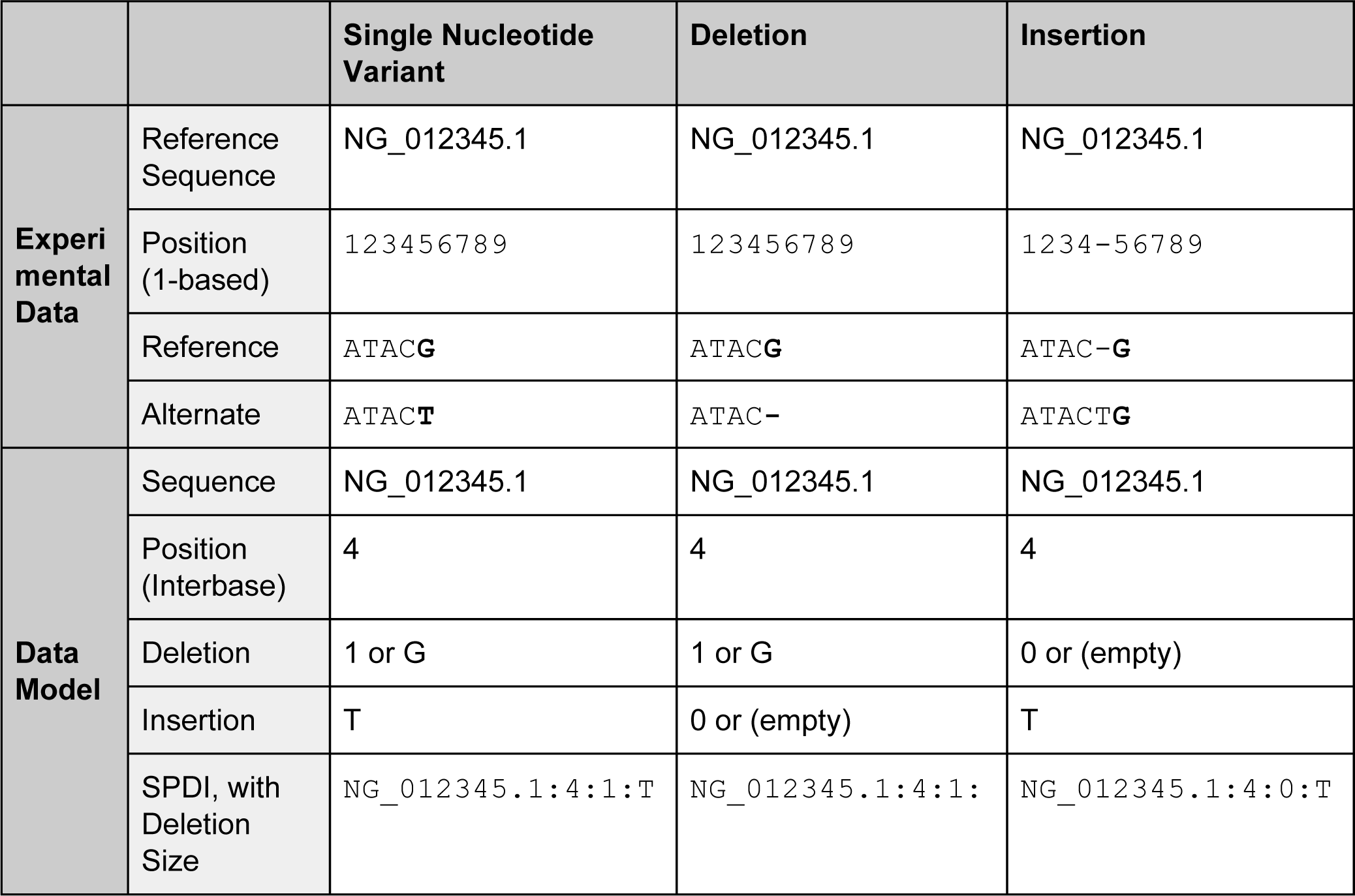
SPDI representation for three simple variant types

|  |  | Single Nucleotide Variant | Deletion | Insertion |
| --- | --- | --- | --- | --- |
| **Experimental Data** | Reference Sequence | NG_012345.1 | NG_012345.1 | NG_012345.1 |
|  | Position (1-based) | 123456789 | 123456789 | 1234-56789 |
|  | Reference | ATAC**G**ACTG | ATAC**G**ACTG | ATAC-**G**ACTG |
|  | Alternate | ATAC**T**ACTG | ATAC-**A**CTG | ATA**C**T**G**ACTG |
| **Data Model** | Sequence | NG_012345.1 | NG_012345.1 | NG_012345.1 |
|  | Position (Interbase) | 4 | 4 | 4 |
|  | Deletion | 1 or G | 1 or G | 0 or (empty) |
|  | Insertion | T | 0 or (empty) | T |
|  | SPDI, with Deletion Size | NG_012345.1:4:1:T | NG_012345.1:4:1: | NG_012345.1:4:0:T |

1. (S)equence: (a string): reference sequence identifier as accession and version,
2. (P)osition: (a non-negative integer) the number of nucleotides to advance on the reference sequence from the boundary before the first nucleotide, which can be thought of as a 0-based interbase coordinate for the variant start,
3. (D)eletion: (a non-negative integer) deletion length, *i.e.* the reference allele length
4. (I)nsertion: (a string) the inserted variant sequence.

SPDI also supports an alternate representation in which the Deletion field is a string containing the literal sequence to delete. This is redundant information but, for efficiency, it facilitates the transformation of contextual alleles to VCF and HGVS without having to refer back to the reference sequences. It also enables some error detection by our services for strandedness and off-by-one errors.

### Representing variants using SPDI

SPDI can represent all variants with defined breakpoints that specify all altered nucleotides. The format is both human- and computer-friendly, namely 4 values separated by colons (Table 1). Notably, the stop coordinate is not stored explicitly, it is implicit from the start location plus the length of the deletion sequence. The explicit start plus the implied stop coordinate are equivalent to the zero-based, half-open interval coordinates used by UCSC genomic resources^2^. By modeling in boundary coordinates, we are able to address precisely insertions as zero-deletion-length variants, which is not possible in a nucleotide coordinate system.

### Terminology: Contextual and Canonical

Services based on the SPDI data model support two kinds of normalization with different properties: the Contextual Allele and the Canonical Allele.

The Contextual Allele is the representation of the variant on a defined reference sequence after correcting for overprecision in low-complexity regions. (Overprecision is defined below.) This reference sequence can be of any type: transcript, protein, or genomic. Because it is based only on the variant and the reference sequence, the Contextual Allele is stable over time. However, it is unique only within a sequence.

The second identifier, the Canonical Allele, extends identification across related sequences. For example, when the A allele of a T>A variant on an earlier sequence version (ie. Chr1 on GRCh37) is later identified as the reference allele in a newer sequence version (ie. Chr1 on GRCh38), that variant is written A= (for reference), and now T is the variant allele, on the new sequence and has a position that differs by thousands of nucleotides from the original sequence. Despite this, they are the same variant because they result in the same local sequence in an equivalent region by alignment. The Canonical Allele is a cluster of all such equivalent contextual alleles. One contextual representation is deterministically chosen as a Canonical Allele Representative and we use its Contextual SPDI as the identifier for the Canonical Allele. Since there is a one-to-one correspondence between the Canonical Allele Representative and the Canonical Allele, we frequently use the term Canonical Allele to refer to both. Each Canonical Allele Representative is a Contextual Allele. The Canonical Allele depends strongly on which regions of which sequences are considered equivalent. Since the sequence alignment can change over time with an improved algorithm and alignment tools, we encapsulate those equivalence interpretations in an Alignment Data Set (ADS) that is versioned Though obvious equivalences are stable, difficult alignments such as those in low-complexity or paralogous regions may be improved. Thus some Canonical Allele Representatives may differ as new ADS versions are released.

### Correcting Overprecision in Variant Assertions

A common problem in variant normalization is that there are multiple ways to describe the same change on the same sequence in the same low complexity region, e.g. homopolymer stretches or tandem repeats. This is termed overprecision for position reporting. Table 2 gives an example of overprecise reporting of variants. Note that the two deletion alleles are the same observation of *ATGACT*, but there are two equally concise ways to represent them against the reference sequence ATGGACT. They are each overly precise, as the variant sequence could have resulted from the loss of the G at either the third or fourth position for the tandem G’s. Likewise, for the overprecision reporting in the insertion example of Table 2.

**Table 2:**
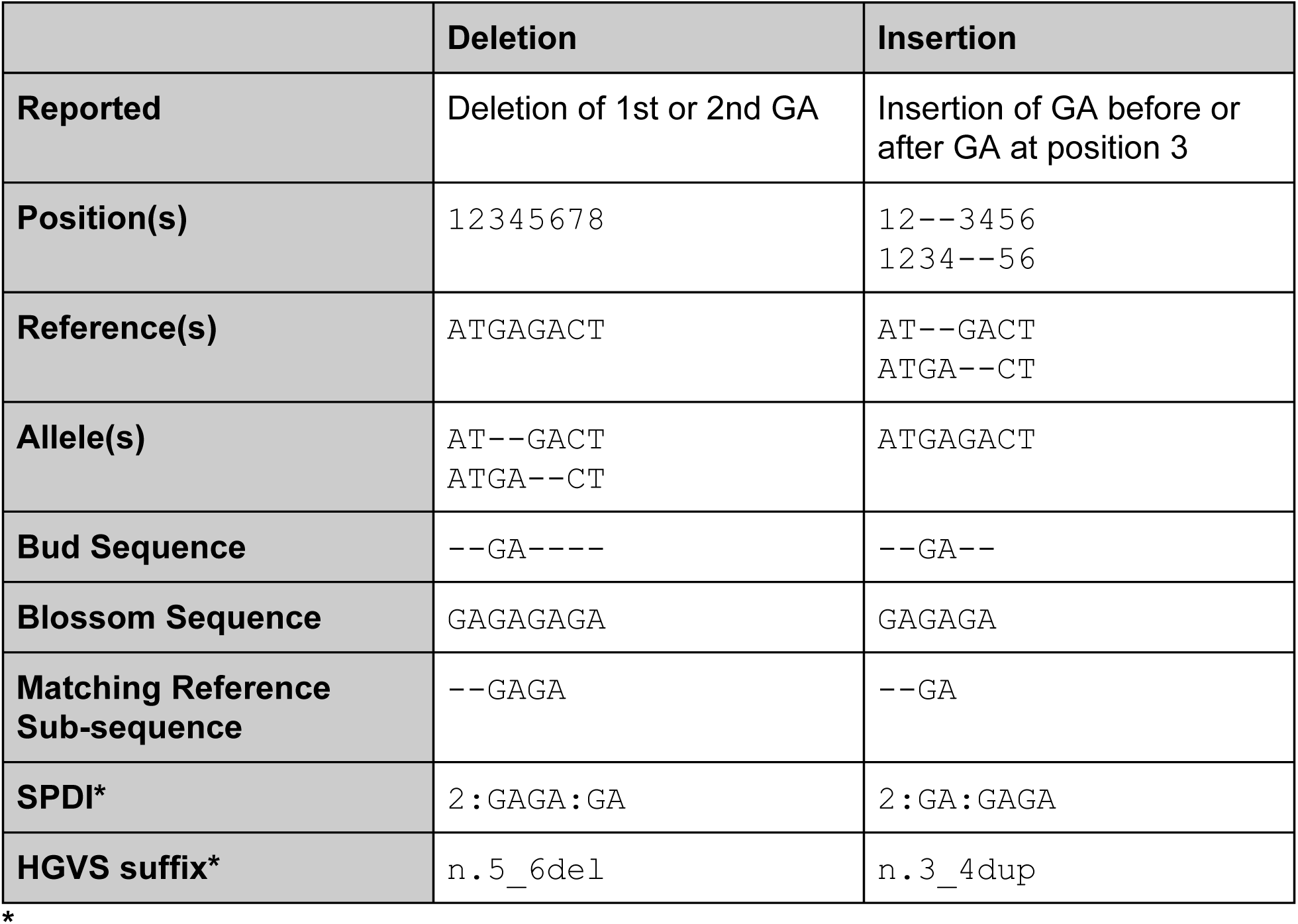
Summary examples of overprecision due to the ambiguity of deleted or inserted nucleotides.

To correct for overprecision, the SPDI services expand submitted variants by applying the NCBI Variant Overprecision Correction Algorithm. Once overprecision is removed, a comparison of two variants asserted on the same sequence model is a straightforward computation using the SPDI format: compare the position, deleted-length, and inserted-sequence. The NCBI Variant Overprecision Correction Algorithm maps observed, non-identity variants to their corresponding Contextual Alleles. It has two phases: shrinking and growing.

### Shrinking

The shrinking phase acts on the deleted and inserted sequences. First, it trims the 3’ ends, one nucleotide at a time, until the two sequences end differently. Next, it trims the 5’ ends (which changes the “from” coordinate in the SPDI model). This is equivalent to “left-shifting” the variation. The resulting sequences have no common nucleotide suffix or -prefix. At least one of the inserted or deleted sequences must be non-empty. Only an identity variant, same as the reference sequence, would produce two empty sequences after shrinking. After the shrinking phase, if neither the deleted sequence nor the inserted sequence is empty, then the variation is not a pure deletion or insertion and so it cannot be overprecise. In these cases, the sequence is now of minimal length, and the algorithm is complete.

### Growing

For pure insertions or deletions, the algorithm chooses the non-empty sequence (either the deletion or insertion sequence) as the bud sequence. Then, it creates a blossom sequence of the same length as the reference sequence by duplicating the bud sequence starting at the “from” position. So, the bud sequence “**GA**” at position 3-4 on a length 8 reference would blossom into “GAGAGAGA". Next, the algorithm chooses the largest interval on the reference sequence matching the blossom sequence and containing the “from” position. This is now the new deleted sequence. Finally, it applies the shrunken variation to this new deleted sequence resulting in the newly inserted sequence. Note, that in the actual implementation we do not materialize the entire blossom sequence. We create it virtually, using a counter-rotating through the bud sequence.

### Remapping (AKA Lift-Over)

Remapping (or lifting over) is a process for calculating equivalent coordinates across sequences using alignments, whether it is between different versions of the same sequence accession, or across sequence types (genomic *vs*. mRNA). NCBI Variation Remapping Service (https://www.ncbi.nlm.nih.gov/variation/services/remapping/) was originally based on the existing assembly-assembly NCBI Genome Remapping Service (https://www.ncbi.nlm.nih.gov/genome/tools/remap) but now includes additional paired alignment sets to provide support for robust remapping and annotation of variants on different sequences. The alignment data set (ADS) includes multiple types of pairwise alignments generated by various NCBI processes including NCBI Splign [9] for cDNA-to-Genomic, NG aligner and NCBI’s assembly-assembly aligner:

- Old assembly to current Genome Reference Consortium (GRC) [10] primary assemblies (e.g. GRCh36(hg17) or GRCh37(hg18) with GRCh38(hg19))
- Patches, alternative loci, or pseudoautosomal regions (PAR) to GRC primary assembly
- RefSeq [11] and select GenBank [12] transcripts to selected RefSeq genomic regions, also known as RefSeqGene (NG), a member of the Locus Reference Genome (LRG) collaboration [13].
- Current RefSeq transcripts (NM/NR/XM/XR) and RefSeq genomic regions (NG) to the latest Assembly
- Previous versions of NG and RefSeq transcripts (NM/NR) to GRC primary assembly

### Alignment Datasets encode Sequence Relationships

Calculating the Canonical Allele from Contextual alleles based on different reference sequences requires a set of invertible relationships between sequences in the ADS. The current ADS covers most scenarios for reporting variants, based on the billions of variants collected by NCBI dbSNP and ClinVar from thousands of different submission sources. However, additional alignment pairs can be created for new reference sequences and added to ADS should the need arise.

The ADS currently consists of nearly 400 thousand pairwise alignments generated from over 300 thousand distinct input sequences. It is stored as binary seq-align objects, consuming 187MB of disk space when compressed. It is regularly updated with new alignments and improved alignment heuristics and software updates. Since the alignments in the ADS encode the sequence relationships, it follows that the Canonical Alleles that depend on those alignments are also regularly recomputed.

Figure 1 shows the pairwise alignments used for remapping an SNV (dbSNP:rs756655831 (www.ncbi.nlm.nih.gov/snp/rs756655831)). The ADS heuristic assumes that alignment is transitive between 1) the sequence and its different versions and 2) between the sequences, NG and NM, that are part of the annotation set for a particular assembly version. Thus, the number of sequence pairs to map between sequences is minimized. Alignment connects the positions on the older sequence models and assembled chromosome sequences through other, more recent, sequences (NM_003193.4, for example).

**Figure 1:**
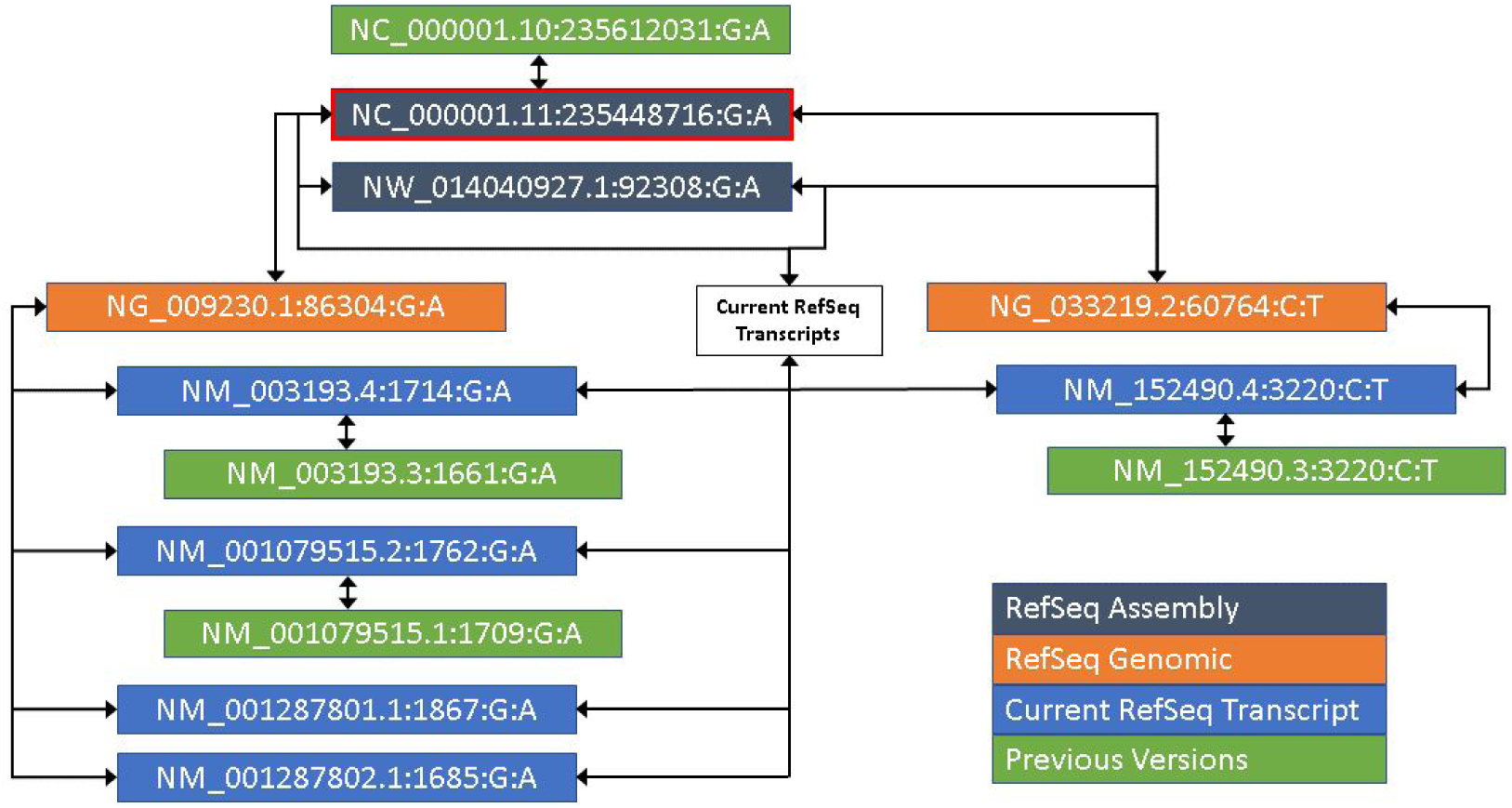
For rs756655831, a representation of the alignments between various sequences, and the resulting SPDIs. Notably, this RefSNP maps to two RefSeqGenes (TBCE NG_009230.1 and B3GALNT2, NG_033219.2). Each has its own set of transcripts, of which all current ones align to the GRCh38 chromosomal sequence NC_000001.11. All previous versions are only aligned to the current versions of the sequence. NW_014040927.1 is a novel patch to chromosome 1 and aligns to only one RefSeqGene, but all transcripts. The location with the red outline is the canonical representation of this set of variants, which allow submissions on any of the SeqIds to be grouped together.

### Alignment Mapping Special Cases

Two special cases must be considered carefully when mapping through alignments: mapping with changed orientation and gaps in the alignment. The forward orientation of mRNA is determined by the direction of transcription (5’ to 3’). As such, the alignments of those genes and their transcript products can be reverse to the chromosome. When mapping variants across such alignments, it is important to reverse-complement the inserted sequence, as the SPDI model uses only the positive strand. In addition, extra care must be taken when mapping an insertion variant with zero-length deletion sequence. Because alignments use base coordinates, not SPDI’s interbase coordinates. It is usually convenient to convert to an insert-before semantic when doing the mapping. However, because directionality changes when strand orientation changes, insert-before becomes insert-after. In order to return to the insert-before semantic, the position must be increased by 1 (Figure 2A).

**Figure 2:**
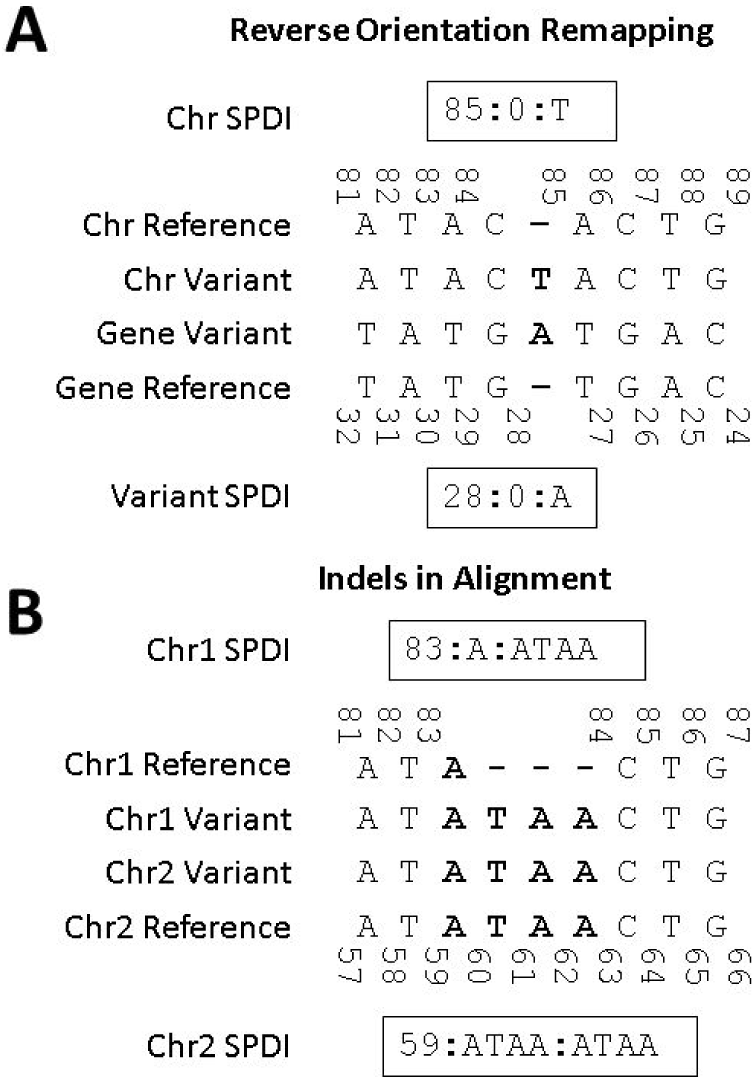
Examples of (A) reverse orientation and (B) indels in alignments special cases. Coordinates are 0-based interbase coordinates. A) For reverse orientation, boundary 85 of the chromosome corresponds to boundary 28 of the gene. But, nucleotide 85 is aligned to nucleotide 27. This must be incremented by one in order to adjust for the change in orientation. (B) In some cases, indels exist in one sequence, but not another. Remapping just the interval may not return any result, as it remaps into a gap. In this example, Chr1 and Chr2 refer to sequential versions of the same chromosome sequence, not two different chromosomes.

In the second case, a disagreement between sequence models may represent an indel variant that is absent in one sequence model and present in another (Figure 2B). Describing the variant represented by these two sequence models results in the variant being described as the *reference* on one sequence model, and an insertion or deletion, or indel, when ambiguities exist, on the other sequence. In general, remapping to other sequences can completely change the type of the variant, between any of the six variant types represented by dbSNP.

### NCBI Variation Services

We implemented public-access API services for the solution presented in this document. It is accessible at http://api.ncbi.nlm.nih.gov/variation/v0.

## Results

SPDI was developed to meet the need of NCBI to represent variants consistently and accurately across resources including dbSNP and ClinVar and for broader use by the community as a web service. dbSNP and ClinVar have tested the SPDI data model and associated algorithms thoroughly and incorporated it into their respective workflows. In so doing, variant aggregation has improved normalization of identical observations submitted with different representations, and to report errors during submission.

### ClinVar

A major function of ClinVar is the aggregation of data from diverse submitters so that each submitter, and the community at large, can determine whether or not there is a consensus in understanding the clinical significance of any allele. That function requires robust, immediate normalization of submissions that may define an allele by non-standard HGVS, by standard HGVS (*e.g.* right-justified, with duplication having precedence over insert), by VCF, or by representation on current or previous versions of any of several reference sequences. From its beginning, ClinVar converted all submissions to HGVS and normalized by determining the corresponding HGVS expression on the reference assembly, currently GRCh38. That HGVS conversion, however, did not standardize the HGVS first, so that if one submitter reported an allele as an insert left-justified and another reported as duplication and another reported as an insert right-justified, identification as the same allele might be missed. As SPDI was adopted into ClinVar’s data flow, these cases could be identified easily. Retrospective correction of alleles by application of the NCBI Variant Overprecision Correction Algorithm has resulted in 1,400 alleles identified as duplicates and merged (Table 3). The majority (about 1,200) are assessed as pathogenic, such as the well-studied 5/7/9T alleles [14] in intron 9 of CFTR that may affect pathogenicity. ClinVar received diverse submitted variant descriptions (Table 4) all of which resolve to the same variant, NM_000492.3(CFTR):c.1210-12T[9] (https://www.ncbi.nlm.nih.gov/clinvar/variation/161188/).

**Table 3:** Summary of ClinVar allele merges after the adoption of the NCBI Variant Overprecision Correction Algorithm.

| Variant Type | Number of merged ClinVar allele |
| --- | --- |
| Total | 1,400 |
| Deletion | 761 (54.4%) |
| Insertion/duplication | 604 (43.1%) |
| Indel | 35 (2.5%) |

**Table 4:**
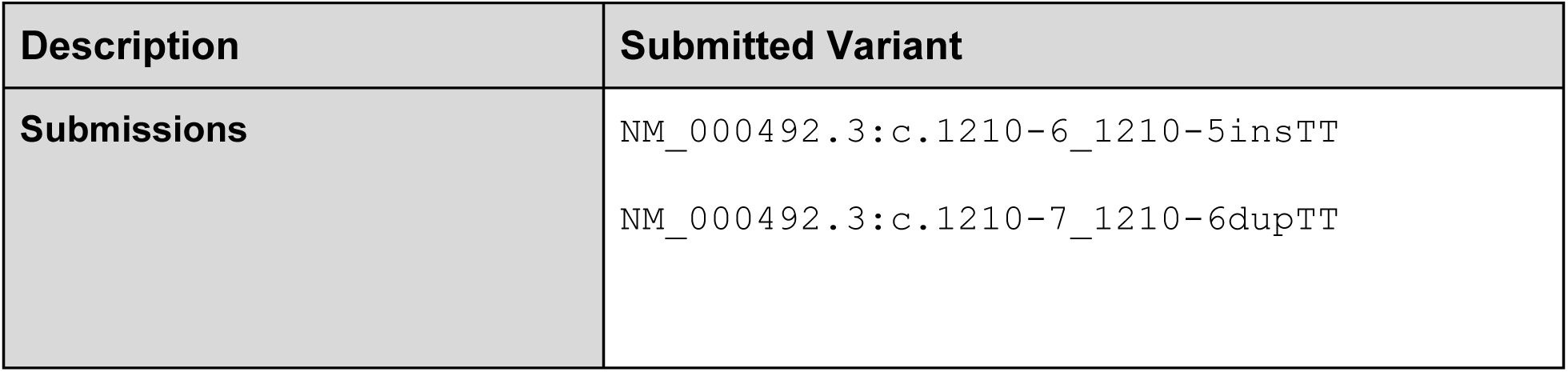
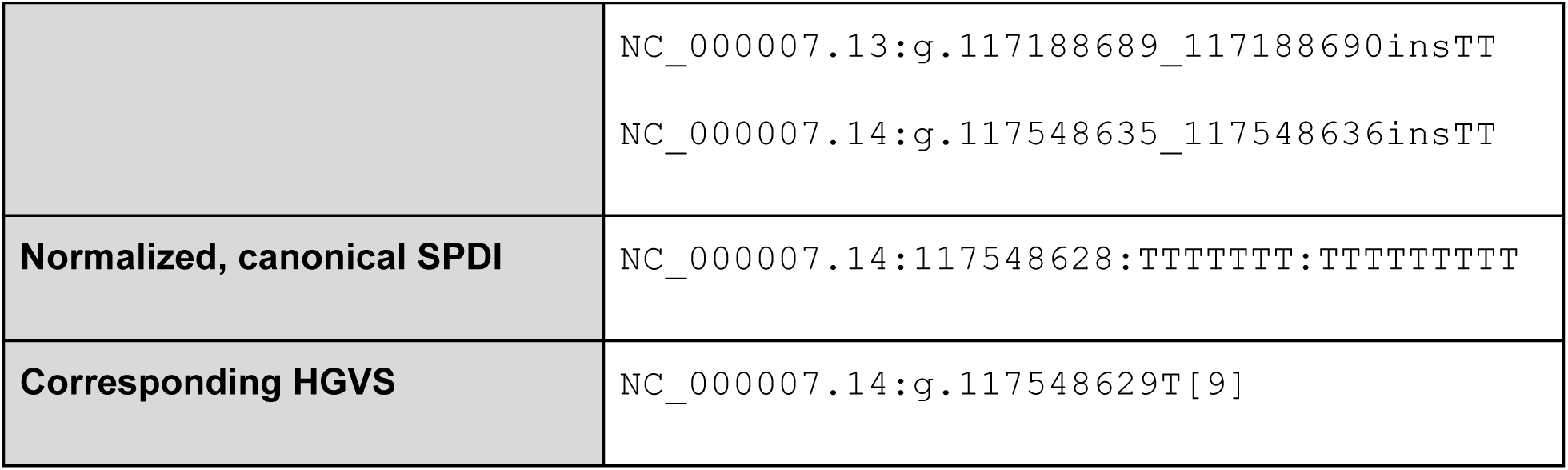
ClinVar submissions for the same allele

| Description | Submitted Variant |
| --- | --- |
| Submissions | NM_000492.3:c.1210-6_1210-5insTT<br>NM_000492.3:c.1210-7_1210-6dupTT |
|  | NC_000007.13:g.117188689_117188690insTT<br>NC_000007.14:g.117548635_117548636insTT |
| **Normalized, canonical SPDI** | NC_000007.14:117548628:TTTTTTT:TTTTTTTTT |
| **Corresponding HGVS** | NC_000007.14:g.117548629T[9] |

### dbSNP

Over the past few years, dbSNP has undergone a significant redesign of its data aggregation and product generation process. For over 15 years, an RDBMS-based solution has served dbSNP well. But with the desire to apply increasingly complex algorithms to identify identical variants and manage over 2 billion human variation submissions (ss IDs), a new system was designed and implemented. The new aggregation and product build pipeline is based on the MapReduce framework with SPDI data model and the correction of over-precision at its core. A set of submitted variants (ss) are aggregated together to form a reference SNP (RefSNP) cluster if they meet two conditions when remapped to a common genomic sequence: on the same deletion interval and same type. dbSNP recognizes six types: identity (observed variants matches the reference sequence), single nucleotide variation (SNV), multiple nucleotide variant (MNV), deletion, insertion, and small insertion and deletion (indel).

All variants in low-complexity regions are now modeled as indels, with the span of the deletion representing the maximum level of precision allowed. This has the particular effect of grouping alleles of varying deletion size in low-complexity regions into a single RefSNP cluster (Figure 3). When applied to dbSNP, 6,423,296 deletions and 2,341,877 insertion variants merged with already existing records (Table 5). All told, 8,945,252 variants merged into 4,493,144 extant RefSNPs, nearly all receiving just one RefSNP. In the most extreme case, one extant RefSNP received 42 merged RefSNPs, rs55883101 (Figure 3). In this particular example, a few of the merged RefSNPs were the result of collapsing of identical alleles (see alleles marked as * and † in 8A). For most of the alleles, it is a variable number of deletions of A were collapsed into a single RefSNP, that now has 43 unique alleles. Notably, rs869211356 did not join this cluster, as its deletion allele begins with a “T”, firmly anchoring the subsequence that is removed. This deletion is precise and uncorrected by the SPDI algorithm. The same is true for all of the SNVs in the region (red boxes).

**Figure 3:**
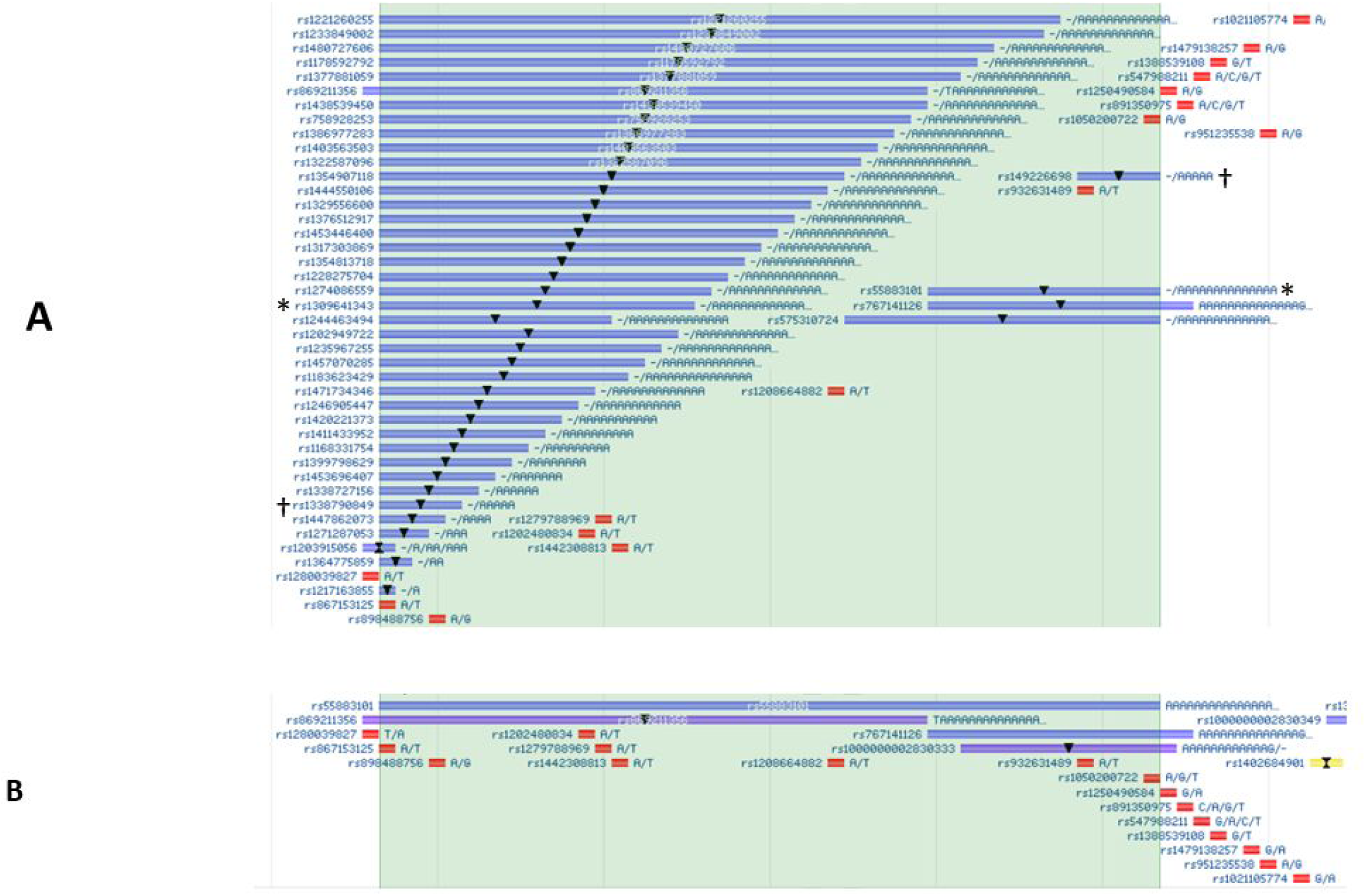
**A)** Example set of RefSNPs that were merged into one (<u>rs55883101</u>) from the previous RDBMS based on build and B) the new distributed dbSNP build that uses SPDI and NCBI Variant Overprecision Correction algorithm. The SNP track coloring scheme has been updated between the two tracks, but in both cases, SNVs are red. In A, deletions are blue with a downward triangle, in B, purple, with a downward triangle. * and † marked variants in A are identical alleles.

**Table 5:** Summary of dbSNP Allele merges after the adoption of the NCBI Variant Overprecision Correction Algorithm.

|  | **Number of Merged RefSNPs** | **Percent of all Merges** | **Percent change of RefSNPs of this Type** |
| --- | --- | --- | --- |
| **All Types** | 8,945,252 | 100% | -1.35% |
| **Deletion** | 6,423,296 | 71.8% | -36.80% |
| **Insertion/Duplication** | 2,341,877 | 26.2% | -42.24% |
| **Indel** | 72,587 | 0.75% | -0.18% |

## Discussion

In their archival function, dbSNP and Clinvar databases accept and store submissions that were ascertained by different projects using different sequencing technologies and variant calling pipelines. Hence the submitted variants can be asserted on different assembly and sequence versions, including older sequence versions from the past 20 years in dbSNP, and represented in different formats including the common HGVS and VCF formats. In addition, there are often redundant variants submitted across submitters and projects that need to be normalized. The databases process the heterogeneous submitted variants and provide non-redundant variant (RefSNP) annotation and reporting on the latest sequence version. This is critical to provide the most accurate view of the variant in sequence context and for efficient data exchange, integration, and reporting. The processing requires mapping all submitted variants, whether defined on genomic or spliced sequences, to the latest assembly version (i.e. GRCh38). Then they can be compared to existing variants in dbSNP and ClinVar to determine if they are novel or not. This can be computationally challenging, which was particularly true for dbSNP that was using a pipeline based on SQL databases and BLAST technologies that weren’t efficient or scalable for processing billions of records. Therefore we developed the SPDI data model to provide a robust and consistent representation of sequence variants. The model supports pipelines for accurate and consistent processing and annotating variants at NCBI, using modern NoSQL computing framework and process control such as Hadoop and Airflow, that is scalable and can be integrated with other NCBI resources. The processes based on SPDI include:

- Validate submitted variant allele and position and convert from HGVS and asserted location (VCF) format to standard Contextual Alleles
- Map variants to NCBI standard top-level sequence on latest genomic assembly version using ADS
- Retrieve all equivalent Contextual Alleles on mRNA, protein, and genomic sequences.
- Obtain equivalent Canonical Alleles for normalizing variants across disparate submissions and to existing reference variant (*i.e*. dbSNP rs).
- Convert variant representations to standard VCF format for export and data exchange

We also made these functions open-access as API calls for users to analyze and normalize their variants that are consistent with NCBI.

### SPDI compared to HGVS

SPDI does not require that the submitter specify a class of variant in its notation; for SPDI, all variants are defined by simple operations of deletion then insertion. HGVS notation, however, requires that when an allele could be described in more than one way, a decision be made as to how a variant is named, according to the priority of “(1) deletion, (2) inversion, (3) duplication, (4) conversion, (5) insertion” (http://varnomen.hgvs.org/recommendations/general/). These priorities are not always followed, so NCBI often receives submissions for inversions as deletion/insertions or for duplications as insertions. Note this list of priorities does not include repeat regions, so there is even more variability within the community for describing alleles such as the polymorphism in intron 9 of the CFTR gene. The standard is to report as a repeat (http://varnomen.hgvs.org/recommendations/DNA/variant/repeated/), yet most submissions are received with HGVS for insertion or a duplication (Table 4). In addition, HGVS requires variants are right-shifted, but again, submissions are received in a variety of shifted states. With SPDI notation there is no interpretation of the type of allele and it is algorithmically unambiguous to generate a standard representation without maintaining complex parsing logic. SPDI currently supports a subset of the variants that can be represented by HGVS nomenclature (see *SPDI limitations* below).

### SPDI compared to VMC

Concurrently during SPDI development, the Global Alliance for Genomics and Health (GA4GH) developed the Variation Modelling Collaboration (VMC) data model for describing variant (manuscript in preparation). VMC and SPDI are similar in that: 1) Both provide specifications for a reference sequence, precise location, and alleles. 2) The VMC “Allele” entity is very similar to SPDI. Both use interbase coordinates and an interval on a reference sequence replaced by a precise sequence specified in IUPAC notation. 3) The variant type is not tagged or labeled but inferred from the variant data in the models. The differences are: 1) The VMC model is broader in scope than SPDI. In addition to representing individual precise variants, VMC also represents precise haplotypes and genotypes. (2) Normalization is required in VMC but optional in SPDI. This allows SPDI to represent over-precise assertions, as they are submitted to dbSNP and ClinVar, that go beyond sequence identity. (3) SPDI uses the NCBI Variant Overprecision Correction algorithm for normalization whereas VMC uses Variant Tool (right shifting)^3^. This means VMC saves space for long variants but the reference interval does not cover all bases that may be affected by the variant. (4) Most operations on VMC require access to the full reference sequence database because VMC alleles specify the interval to be replaced via an identifier. SPDI can encode the actual reference allele sequence in the variant. (5) Data transformation from SPDI to different formats (i. e. HGVS or VCF) is easier as a consequence of (2) and (3). A normalized SPDI encodes the residues of the reference sequence, it is possible to generate normalized forms according to other algorithms: whether left-shifting or right shifting. (6) Although limited in scope, SPDI variant representation is shorter and easier to digest as a single human-readable line containing the sequence, position, and the actual allele sequences and yet machine parseable for developing parallelize workflow without the need for retrieving the reference sequence. VMC is intended for machine-machine communication, so has a heavily nested structure filled with long, random identifiers.

### Variation services compared to existing tools and services

SPDI and the associated NCBI Variation Services have some functions that are similar to existing tools and services including ClinGen Allele Registry [15], Variant Validator [16], Ensembl Variant Recoder, TransVar [17], Mutalyzer [18], VEP [19], and snpEff [20]

(Table 6). Of all the existing tools, the NCBI Variation Services are most similar in functions to the ClinGen Allele Registry that provide functions such as variant ID search, remapping, and normalization. Both services accept common input formats such as HGVS, VCF, and dbSNP RS and provide JSON output format that is amenable to computational processing. A distinguishing feature of SPDI is it provides original annotations and supporting variant submission evidence along with frequency data and publications for over 670 Million variants that exist in dbSNP [2] and ClinVar [3]). The Allele Registry provide links to the annotations and dependent on users to register the variant or data releases from primary sources such as dbSNP or ClinVar. SPDI services integrate dbSNP, ClinVar, and other NCBI resources such as Assembly and RefSeq to provide timely and regular updated when new data are available. Variants are mapped to the latest genomic and RefSeq mRNA and protein sequences and annotated with functional consequences, allele frequency, publications (PubMed), and ClinVar clinical significance. Using alignment data sets that include more than 400,000 paired alignments the service can provide remapping across more than 300,000 unique sequences.

**Table 6:** Comparison of NCBI Variation Services to other common variant analysis tools.

| Tools and Services | Input Format | Output Format | Variant Normalization | Included annotation source | Reference sequence source | Remapping |
| --- | --- | --- | --- | --- | --- | --- |
| NCBI Variation Services (SPDI) | SPDI, HGVS, VCF, dbSNP RS | JSON | Yes | dbSNP; ClinVar | GenBank, RefSeq | Yes |
| ClinGen Allele Registry<sup>4</sup> | HGVS, dbSNP RS, and many local identifiers | JSON | Yes | Linkouts to various sources | GenBank, RefSeq, Ensembl, and others | Yes |
| Variant Validator<sup>5</sup> | HGVS, VCF | Tab-delimited, JSON<sup>6</sup> | Yes | dbSNP; ClinVar | RefSeq, LRG | Yes |
| Ensembl Variant Recoder<sup>7</sup> | variant identifier, HGVS | JSON, XML | No | dbSNP, EBI | Ensembl | No |
| TransVar<sup>8</sup> | HGVS | Tab-delimited | No | None | Ensembl, RefSeq | No |
| Mutalyzer<sup>9</sup> | HGVS, dbSNP RS | Tab-delimited | No | None | RefSeq | Yes |
| VEP<sup>10</sup> | HGVS, dbSNP, COSMIC, HGMD identifiers, VCF | JSON, XML, VCF | No | dbSNP, ClinVar, EBI, COSMIC, and Others | Ensembl, RefSeq | Yes* |
| snpEff<sup>11</sup> | VCF | VCF, HGVS | No | Various Public Databases | Various Public Databases | No |

\*VEP supports remapping between transcripts, and transcripts to genomic, but not between different genomic sequences.
<sup>4</sup> http://reg.clinicalgenome.org/. Accessed 16 Aug. 2018. (PubMed: 30311374).
<sup>5</sup> https://variantvalidator.org/. Accessed 16 Aug. 2018. (PubMed: 28967166).
<sup>6</sup> https://rest.variantvalidator.org/webservices/variantvalidator.html. 6 March. 2019.
<sup>7</sup> http://www.ensembl.org/info/docs/variation. Accessed 16 Aug. 2018.

### Limitations of SPDI and associated Variation Services

While SPDI provides a powerful representation of precise sequence changes relative to a reference, there are some limitations.

- SPDI does not support reporting a position offset from the reference such as used in HGVS expressions like c.-7A>C, c.88+2T>G, c.89-1G>T, or c.*23T>C Instead it is recommended to select as reference a sequence that includes the variant in order to reduce computation complexity and ambiguity by sequence features context such as CDS annotation that can change.
- Variants without precise breakpoints (such as large structural variants detected by paired-end mapping) cannot be specified in our model.
- NCBI Variation Services only partially supports variants with a protein reference.
- While the services do compute a contextual allele, they are not remapped as we have no support for codon degeneracy.
- NCBI Variation Services only supports variants reported on reference sequence (RefSeq) which may not represent all common and alternate haplotypes.

### Using the Contextual and Canonical Allele

The Canonical Allele representations are essential for determining if two variants represented on different reference sequences) are the same – yet this determination is an *interpretation* based on the preferred reference sequence and available alignments, and subject to change and refinement over time. As we mentioned in the method section, the precision-corrected Contextual Allele on a particular sequence accession and version is stable over time in dbSNP and ClinVar. It serves as the archival summary evidence for the variant observed. In contrast, the Canonical Allele can change over time because it is computed using the Contextual Allele on the original asserted sequence and alignments that can change with any new sequence version or alignment improvements. Therefore, we propose that whenever the Canonical Allele is used, the Contextual Allele should also be made available to allow users to see supporting observed variant as well as the derived interpretation.

## Conclusion

The SPDI data model and associated variation services were developed to address the challenges of processing, annotating, and exchanging the growing volume of variation data in dbSNP and ClinVar databases. This work has resulted in improving the identification and validation of submissions and standardizing their representation. The APIs are provided as a public service to allow users to interrogate and process their variants based on the SPDI model that is consistent with the usage by dbSNP and ClinVar

## Acknowledgments

We would like to thank the GA4GH GKS Variant Representation members, Sarah Hunt, Raymond Dalgleish and Peter Causey-Freeman for their helpful discussions and feedback.

This work was supported by the Intramural Research Program of the National Library of Medicine, National Institutes of Health.

## Footnotes

1 “All of Us Research Program - NIH.” https://allofus.nih.gov/. Accessed 15 Aug. 2018.

2 “The UCSC Genome Browser Coordinate Counting Systems | UCSC “ 12 Dec. 2016, http://genome.ucsc.edu/blog/the-ucsc-genome-browser-coordinate-counting-systems/. Accessed 9 Jan. 2019.

3 “Vt - Genome Analysis Wiki.” https://genome.sph.umich.edu/wiki/Vt. Accessed 9 Jan. 2019.

4 http://reg.clinicalgenome.org/. Accessed 16 Aug. 2018. (PubMed: <u>30311374</u>).

5 https://variantvalidator.org/. Accessed 16 Aug. 2018. (PubMed: <u>28967166</u>).

6 https://rest.variantvalidator.org/webservices/variantvalidator.html. 6 March. 2019.

7 http://www.ensembl.org/info/docs/variation. Accessed 16 Aug. 2018.

8 http://bioinformatics.mdanderson.org/transvarweb/. Accessed 16 Aug. 2018. (PubMed: 26513549).

9 https://www.mutalyzer.nl/. Accessed 16 Aug. 2018. (PubMed: <u>18000842</u>).

10 https://useast.ensembl.org/info/docs/tools/vep/index.html. Accessed 16 Aug. 2018. (PubMed: 27268795).

11 http://snpeff.sourceforge.net/. Accessed 16 Aug. 2018. (PubMed: <u>22728672</u>).

